# Reduced-Order Modeling and the Physics Governing Flapping Wing Fluid-Structure Interaction

**DOI:** 10.1101/2021.06.11.448136

**Authors:** Ryan Schwab, Erick Johnson, Mark Jankauski

## Abstract

Flapping, flexible insect wings deform during flight from aerodynamic and inertial forces. This deformation is believed to enhance aerodynamic and energetic performance. However, the predictive models used to describe flapping wing fluid-structure interaction (FSI) often rely on high fidelity computational solvers such as computational fluid dynamics (CFD) and finite element analysis (FEA). Such models require lengthy solution times and may obscure the physical insights available to analytical models. In this work, we develop a reduced order model (ROM) of a wing experiencing single-degree-of-freedom flapping. The ROM is based on deformable blade element theory and the assumed mode method. We compare the ROM to a high-fidelity CFD/FEA model and a simple experiment comprised of a mechanical flapper actuating a paper wing. Across a range of flapping-to-natural frequency ratios relevant to flying insects, the ROM predicts wingtip deflection five orders of magnitude faster than the CFD/FEA model. Both models are resolved to predict wingtip deflection within 30% of experimentally measured values. The ROM is then used to identify how the physical forces acting on the wing scale relative to one another. We show that, in addition to inertial and aerodynamic forces, added mass and aerodynamic damping influence wing deformation nontrivially.

## 1. Introduction

Insects leverage flexible, flapping wings to realize flight. Unsteady aerodynamic forces generated by flapping wings allow insects to hover or advance forward [1], while aerodynamic and inertial asymmetries caused by differences in left-right flapping kinematics enable insects to perform complex aerial maneuvers [2]. Through a combination of experimental, analytical and numerical studies, tremendous progress has been made over the past several decades to characterize the aeromechanics of insect flight during hovering [3–5], climbing [6–8] and maneuvering [9–11]. More recently, the wing structure itself has garnered attention, where structural features such as venation, corrugation, and camber have been determined to augment aerodynamic performance [12–15].

During flight, insect wings deform under both aerodynamic and inertial-elastic forces [16]. The fluid and structural physics are tightly coupled, where aerodynamic forces influence how the wing deforms and resulting wing deformation influences the surrounding flow field. Computational methods, typically computational fluid dynamics (CFD) coupled to finite element analysis (FEA) based solvers, have been used to simultaneously resolve the instantaneous wing shape and surrounding flow field [12,17–24]. While direct computational methods can provide accurate estimates of flapping wing dynamics, they often require computational times on the order of hours to days to generate solutions. Both CFD and FEA face computational challenges, and their individual challenges are compounded when the two computational solvers are coupled together. CFD necessitates the Navier-Stokes equations be solved across a discretized fluid domain [25], which may result in tens of thousands of conditions that must be satisfied at each time interval of analysis. From the structural perspective, the large rotational kinematics of flapping wings give rise to centrifugal forces that periodically influence the wing’s stiffness [26]. Centrifugal effects require that the FEA stiffness matrix be updated at each interval of analysis if the angular velocity is not constant. The high computational resources demanded by direct computational methods therefore render them unsuitable for parametric studies that consider variable wing geometry, kinematics or structural properties. Moreover, reliance on computational models may obscure the physical phenomena governing (e.g., aerodynamic drag, added mass, etc.) wing deformation.

In this regard, analytic models may be preferable to computational models because (1) they provide physical insights into flapping wing dynamics, and (2) in some cases, can identify response trends more quickly due to lower solution times. The most commonly used analytic models in flapping wing literature are based on quasi-steady blade element theory (BET) [27]. BET assumes that a wing can be discretized into chordwise strips (blade elements). Based on known flapping kinematics and measured or estimated aerodynamic coefficients, thin airfoil theory is used to predict aerodynamic forces and moments acting on a single blade element, and these differential forces are integrated over the wing surface to estimate total aerodynamic forces. BET is often restricted to describing the aerodynamics of rigid wings, though Wang et al. developed a fluid-structure interaction (FSI) model based on BET to estimate the aerodynamics of twistable wings [28]. Other researchers have used Theoderson’s unsteady aerodynamic model to predict flapping forces of flapping wings [29–31]. Kodali et al. derived an analytical FSI model of a flexible pitching, plunging wing based on Theoderson’s model. Their model predicted wingtip deflection with reasonable accuracy compared to experimental and numerical findings [32]. However, Theodorsen’s model is based on inviscid assumptions, and may be limited in contexts where drag influences wing loading considerably; in these cases, BET may perform more favorably than Theoderson’s model.

Based on this review, the first objective of this research is to develop a reduced-order FSI model of a flapping wing experiencing single-degree-of-freedom (SDOF) rotation. Aerodynamic drag is appreciable when the wing is undergoing SDOF rotation. The fluid model is based on BET, but because it permits elastic deformation, we refer to it as *deformable* blade element theory (DBET). The analytical model provides insights into the physics governing flexible wing aerodynamics and expands upon the research in [33]. The second objective of this work is to validate the reduced-order model against experimental results as well as solutions generated via high-fidelity coupled CFD/FEA. This allows us to quantitatively compare accuracy and solution times achievable by both low and high-order models. While the SDOF flapping kinematics in this work are simplified from the multiple-degree-of-freedom (MDOF) wing kinematics of flying insects, this work is a foundational step towards reduced-order modeling of FSI in wings subject to more realistic motions.

The remainder of this manuscript is organized as follows. First, we derive the reduced-order FSI model based on DBET and the assumed mode method (AMM). Next, we detail the high-fidelity coupled CFD/FEA used to predict wing deformation. We then present a simple experiment to assess the accuracy of both models. We conclude by discussing insights gained from the DBET model and comment on its efficiency relative to the computational model.

## 2. Theory

Here, we present two models capable of resolving the deformation of a wing experiencing SDOF rotation. The first model is low-fidelity and semi-analytical, whereas the second model is high-fidelity and computational. The objective of this research is to compare the accuracy and computation times achievable by both models. Note that the following FSI frameworks are general and applicable to a range of wings and SDOF flapping kinematics; for this reason, the simulation parameters specific to experimental validation are presented in Section 3.

### (a) Reduced-Order Modeling

The reduced-order FSI model is based upon AMM and DBET. The structural and aerodynamic drag models originated in [33], however this previous work did not compare the ROM to high-fidelity coupled CFD/FEA. This research further advances the previous ROM by incorporating added mass, which we show influences the wing’s perceived stiffness and increases aerodynamic loading.

#### (i) Structural Model

The motion of a flapping wing can be modeled as a superposition of elastic deformation on top of larger rigid body rigid body rotation. Rigid body rotation is generally an active or controlled degree-of-freedom, whereas elastic deformation occurs passively under internal and external forces. Provided elastic deformation is small, it can be treated as a summation of vibration mode shapes multiplied by their modal responses, also called modal participation factors. The benefit to formulating the structural model in terms of modal coordinates instead of physical coordinates is that the wing’s mode shapes are independent of dynamic inputs. They can be pre-computed prior to dynamic simulation based on knowledge of the structure alone. Further, we can truncate higher modes that do not contribute meaningfully to wing deformation (e.g., modes that correspond to natural frequencies substantially outside of the range of input frequencies), which reduces the computational time required to solve the structural model.

We establish a reference frame that rotates with the rigid body rotation of the wing (Fig. 1). An inertial *XY Z* coordinate system undergoes a rotation of magnitude *α*, where *α* is the wing’s flapping angle. The resulting *xyz* coordinate system has an angular velocity ***Ω***, where

**Figure 1.**
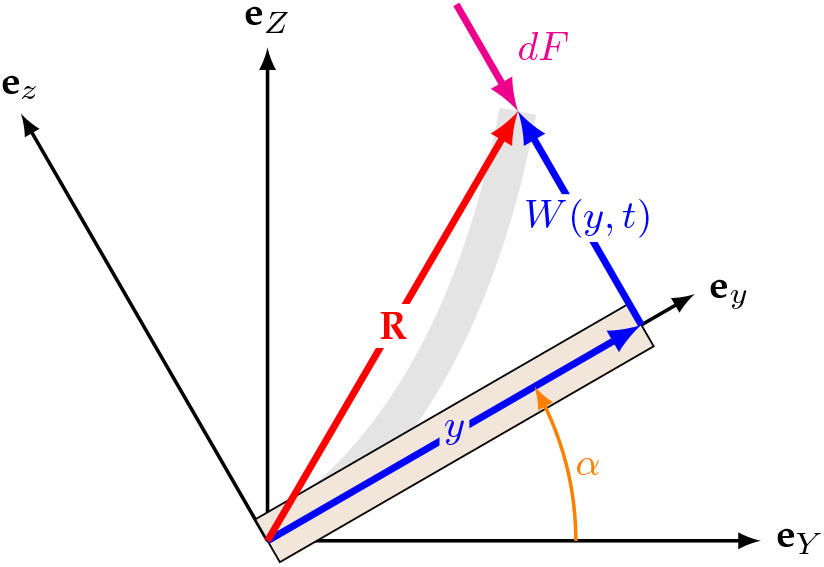
An *xyz* reference frame rotates with the rigid body rotation of the wing, where *α* denotes the wing’s flap angle. Within the rotating reference frame, the wing experiences small out-of-plane deflection *W* (*y, t*), where *y* is the axial location of a point on the wing. The wing is subject to an aerodynamic force per unit area *dF*. Gray region shows a deformed state of the wing.

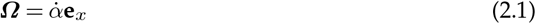

where ·denotes a derivative with respect to time and **e**_*n*_ denotes a unit vector in the n direction. Within the *xyz* frame, we draw a position vector **R** from a fixed point of rotation to a differential mass element *dm* (Fig. 1), where

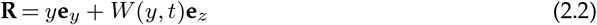

Above, *y* denotes the planar coordinates of the differential mass and *W* (*y, t*) describes and a small out-of-plane elastic deformation. In-plane deformation and twisting about the *y* axis are neglected. The out-of-plane deformation can be expanded via the separation principle such that

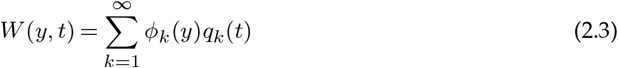

where *ϕ*_*k*_ (*y*) is the wing’s *k*^*th*^ mode shape normalized with respect to the wing’s mass and *q*_*k*_ is the corresponding time-dependence, or modal participation factor to be determined. *ϕ*_*k*_ (*y*) is a static quantity and can be determined either via modal analysis in FEA or analytically for simple structures. The velocity 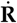 of the differential mass is

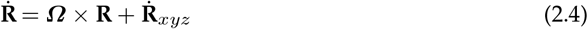

where 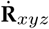 is the differential mass velocity referenced from the rotating frame. The kinetic energy *T* of the entire wing is

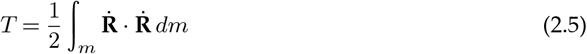

which can be represented in modal coordinates as

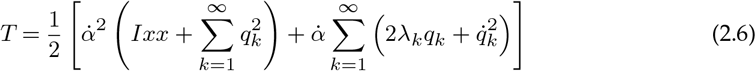

where *I*_*xx*_ is the wing moment of inertia about *x* and *λ*_*k*_ is an inertial constant defined by *λ*_*k*_ *=∫*_*m*_ *yϕ*_*k*_ *dm*. The wing’s potential energy *U* is

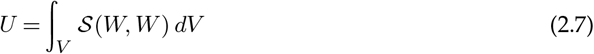

where *𝒮* is a symmetric, quadratic strain energy density function and *V* is the wing’s volume. By applying Lagrange’s equation, we arrive at the equation governing modal response *q*_*k*_ as

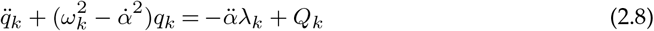

where *ω*_*k*_ is the wing’s *k*^*th*^ natural frequency *in vacuo* and *Q*_*k*_ includes all non-conservative modal forces to be determined. The above equation is linear and time-varying, where the wing stiffness is influenced by its angular rate of rotation through a phenomena called centrifugal softening. Lastly, a general physical force *d***F** per unit area is converted to the *k*^*th*^ modal force by

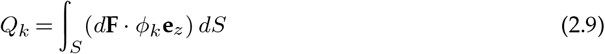

where *S* is the surface over which the force acts. In the following sections, we determine the modal forces *Q*_*k*_ associated with aerodynamic drag and added mass via DBET.

#### (ii) Aerodynamic Drag

Due to simplified SDOF flapping kinematics, the wing will produce negligible lift and drag becomes the dominant aerodynamic force. Assuming drag acts only in the *z* direction and does not vary over the wing’s chord, the drag per unit area is *d***F**_*D*_ is

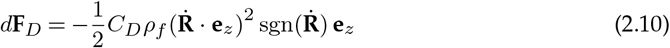

where *C*_*D*_ is an empirical drag coefficient and *ρ*_*f*_ is fluid density. *C*_*D*_ typically varies with respect to the wing’s angle of attack, but in this case can be treated as a constant since the wing’s angle of attack is always *π/*2 or 3*π/*2 because in a static fluid air is only moved normal to the wing surface. Expanding this expression in terms of 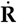 and the expansion in Eq. 2.3 gives

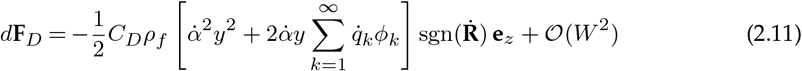

where we neglect terms of *𝒪* (*W*)^2^ because deformation is small. Projecting the physical drag force into the modal domain via Eq. 2.9 yields two modal force terms. The first modal force term is

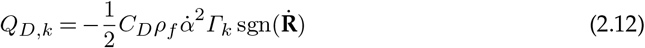

where *Γ*_*k*_ is a constant defined by *Γ*_*k*_ =∫_*y*_ *b*(*y*)*y*^2^ *ϕ*_*k*_ *dy* and *b*(*y*) is the wing chord. *Q*_*D,k*_ depends only on the rigid body rotation of the wing and is not significantly affected (besides the sgn term) by elastic deformation. We therefore refer to *Q*_*D,k*_ as rigid body drag. The second modal force term is

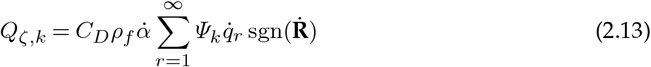

where *Ψ*_*k*_ is a constant defined by *Ψ*_*k*_ =∫_*y*_ *b*(*y*)*yϕ*_*r*_*ϕ*_*k*_ *dy*, and *r* is a new modal index inclusive of *k*. Unlike the aerodynamic loading term, *Q*_*ζ,k*_ is a function of the wing’s elastic deformation velocity as well as its rigid body rotation. It effectively behaves as a time-periodic aerodynamic damping term that attenuates the elastic oscillations of the wing. For this reason, we refer to *Q*_*ζ,k*_ as aerodynamic damping hereafter.

#### (iii) Added Mass

Added mass is a phenomena where a volume of air displaced by an accelerating structure increases the effective inertia of that structure. Insect wings have a low surface density and high surface area, so added mass cannot safely be neglected. In this section, we derive an expression to incorporate added mass into the FSI model.

Added mass is proportional to the differential mass’s out-of-plane acceleration **a**_*z*_ given by

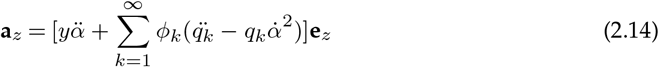

From [34], the added mass force per unit length *d***F**_*am*_ for a thin two-dimensional wing section is

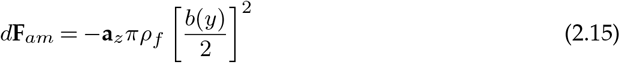

Substituting **a**_*z*_ into the above and converting the physical force to the modal domain through Eq. 2.3 yields three terms. The first added mass term is

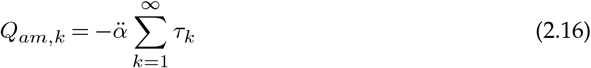

where *τ*_*k*_ = *πρ*_*f*_ ∫ _*y*_[*b*(*y*)*/*2]^2^*yϕ*_*k*_ *dy*. Similar to rigid body drag, this added mass term is a function of rigid body rotation only and is not influenced by the elastic deformation of the wing. We will refer to this term as rigid body added mass. The second added mass term is

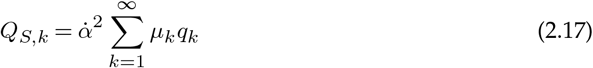

where 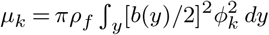. Interestingly, *Q*_*S,k*_ is influenced both by the wing’s angular velocity as well as its instantaneous shape. Similar to the centrifugal softening observed in the structural model, this term modulates the wing’s stiffness periodically; for this reason, we refer to *Q*_*S,k*_ as added mass stiffness. The final added mass term is

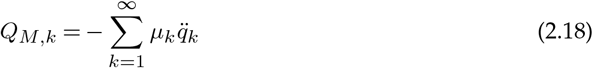

where *Q*_*M,k*_ is proportional to the acceleration resulting from structural deformation. This term adds to the perceived mass of the wing if the wing is vibrating in air, and thus we refer to it as added mass inertia. Due to added mass inertia, the natural frequencies of the wing are lower in air than in vacuum.

### (b) Computational Modeling

Numerical studies focused on flapping wings often rely on CFD and FEA. The accuracy of these direct numerical models is generally assumed to be better than that of their reduced-order counterparts, though at the expense of greater computational costs. Here, we develop a two-way coupled CFD and FEA model to predict the dynamic response of the flapping wing. This high-fidelity computational model enables us to incorporate dynamic phenomena neglected by the DBET model, such as unsteady aerodynamics and structural non-linearity. For our model, we use Siemens’ Star-CCM+ (v15.04.008) CFD package and Dassault Systèmes Abaqus 2019 (6.19-1) FEA package interfaced via Co-simulia.

CFD calculates the pressure on the wing’s surface and surrounding flow field by solving the Navier-Stokes equations. In order to resolve turbulence without requiring untenable computation times, we rely on a Spalart-Allmaras (SA) model to close the Reynolds-averaged Navier-Stokes (RANS) equations. RANS formulations are the most commonly used methods for modeling turbulent flows, and the SA model was chosen for its efficiency as a one-equation model as well as its efficacy in aerodynamic and transient flows. We use a Chimera mesh approach to account for the larger rigid-body rotation of the flapping wing [35]. This method uses multiple meshes (Fig. 2) – one that rotates with the wing (the *overset* mesh) and another that remains stationary and describes the entire fluid domain (the *background* mesh). Interpolation between the boundary of the overset mesh and the background mesh allows the conservation of mass and momentum to be maintained across both regions. In addition to the large prescribed rotations, there is a significant deformation on the wing due to its flexibility. To account for this deformation, our overset mesh region uses a radial basis function (RBF) mesh morphing method to stretch and compress elements around the wing as it deforms. These meshing methods allow the CFD model to run dynamically without the requirement of remeshing the domain at each timestep.

**Figure 2.**
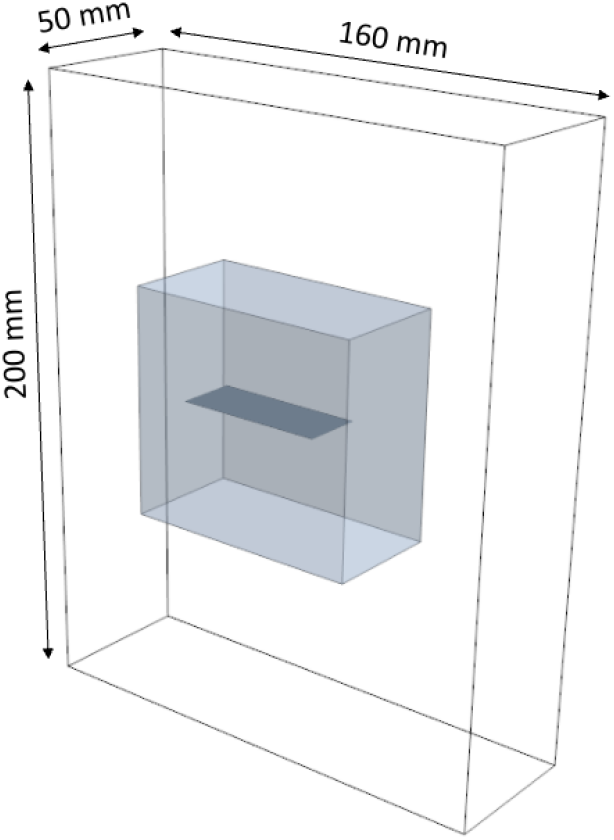
Background mesh (dimensions labeled) and overset mesh (grey) of the fluid domain in Star-CCM+. The rectangular wing is seen in dark grey within the overset mesh. The background mesh must be large enough to accommodate flapping kinematics, while the overset mesh must accommodate the out-of-plane deformation of the wing.

We used Abaqus FEA to resolve wing deformation resulting from aerodynamic loads interpolated onto the wing from the Star-CCM+ simulation and the prescribed rotational kinematics. The FEA model incorporates nonlinear geometric effects to account for the large displacements associated with our prescribed flapping profile. To apply the flapping kinematics to the FEA model, we imposed pinned boundary conditions along the wing’s root edge. The root edge was prescribed a periodic angular displacement consistent with the rotation amplitudes and frequencies described in the simulation parameters section. We use an implicit solver to calculate the spatiotemporal deformation field at each time step.

Lastly, the CFD and FEA models are integrated via Abaqus Co-simulia. This allows for communication between the structural and fluid solvers at each timestep. Abaqus leads the time marching by first applying the prescribed kinematics described above, then resolving the structural deformation of the wing. The wing’s new geometry and rigid body rotation are then sent to Star-CCM+ in terms of its nodal displacements. Star-CCM+ applies the imported displacements to its wing geometry and uses the displacements for the initiation of the RBF morphing of the overset mesh. The fluid domain in Star is resolved based on the impact of the wing’s rotation and deformation on the fluid, and then the shear and normal pressures on the wing surfaces are exported and sent back to Abaqus. Finally, the wing’s structural deformation is resolved in Abaqus, and the process repeats until a quasi-steady state response and desired physical time are reached.

## 3. Experimental Methods

We developed a simple experiment to evaluate the accuracy of the flapping wing FSI models. We used a mechanized rotation stage to prescribe flapping kinematics to a thin paper wing. Wing deformation was recorded via high-speed videography and wingtip displacement was calculated relative to the rigid body rotation of the wing using motion capture.

The mechanized rotation stage (Fig. 3) used for all experiments is summarized in [33]. Flapping trials were filmed with a high-speed video camera (Krontech, Chronos 2.1-HD) at 2996 frames per second with a spatial resolution of 1280 x 512 pixels; see supplementary material for a video of the wing flapping at 10 Hz. The experiment was back-lit using a Godox SL-200W LED studio light. Recordings were post-processed in MATLAB. The tip of the flexible wing and the reference plate were tracked in a world frame using the DLTdv digitizing tool from the Hedrick lab [36]. We placed a thick metal plate behind the flexible paper wing to serve as a reference for the wing’s rigid body. The rotation angle was measured from the center of rotation and the tip of the reference plate. The out-of-plane displacement of the flexible wing was determined with respect to a rotating coordinate system attached to the reference plate. We identified wingtip displacement for three consecutive flapping periods and then fit the wingtip displacement as a function of time using a fifth order Fourier series. We compared curve-fitted wingtip displacement to model predictions.

**Figure 3.**
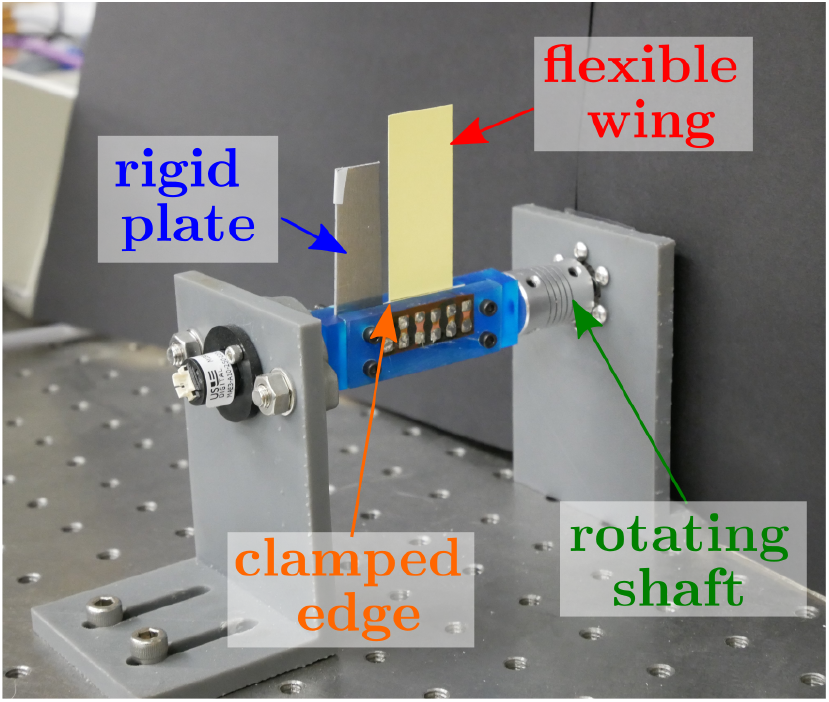
Mechanized SDOF rotation stage used to drive the rigid body rotation of the flexible wing.

### (a) Experimental & Simulation Parameters

All simulation properties are summarized in Tab. 1. The paper wing’s length, chord width and surface area are similar to that of a Hawkmoth *Manduca sexta* forewing, though the paper wing weighs about four or five times as much; about 200 mg for the paper wing compared to 40 mg for the moth wing [37]. *M. sexta* are a common model organism in the study of flapping wing flight.

**Table 1.** Simulation parameters.

| Wing Parameters |  |  |  |
| --- | --- | --- | --- |
| Variable | Description | Value | Unit |
| $L$ | Length | 5 | cm |
| $w$ | Width | 2 | cm |
| $t$ | Thickness | 0.017 | cm |
| $\rho$ | Density | 1.2 | g/cm<sup>3</sup> |
| $E_{DBET}$ | Young's modulus (DBET) | 9.5 | GPa |
| $E_{COMP}$ | Young's modulus (Computational) | 9.085 | GPa |
| $f$ | Flap frequency | 8 - 12 | Hz |
| $\alpha_0$ | Flap angle | 0.9-1.01 | Rad |
| Fluid Models |  |  |  |
| Variable | Description | Value | Unit |
| $\Delta t$ | Time step | 1 | ms |
| $\rho_f$ | Air Density | 0.001250 | g/cm<sup>3</sup> |
| $C_D$ | Drag Coefficient | 3.4 | - |
| $\mu_a$ | Dynamic Viscosity of Air | $1.855 \times 10^5$ | Pa-s |
| | Dimensions of the background mesh (L × W × H) | 160 X 50 X 200 | mm |
|  | Mean element length of the background mesh | 1.2 | mm |
|  | Number of elements in the background mesh | 641,817 | - |
| | Dimensions of the overset mesh (L × W × H) | 40 X 80 X 60 | mm |
|  | Mean element length of the overset mesh | 0.6 | mm |
|  | Mean element length of the wing surface | 0.6 | mm |
|  | Number of prism layers off the wing | 5 | - |
|  | Prism-layer growth rate | 1.5 | - |
|  | Number of elements in the overset mesh | 535,653 | - |

Experimental studies show that *M. sexta* flap at about 1/3 the natural frequency of their forewing [38], and research suggests this flapping-to-natural frequency ratio is aerodynamically and energetically beneficial as well [39–42]. Consequently, we wanted to ensure experiments encompassed the 1/3 flapping-to-natural frequency ratio. We measured the first natural frequency of the wing by displacing it and allowing it to freely vibrate. Free vibration was recorded using a laser vibrometer (Polytec, PSV-400). The first natural frequency was 30.1 Hz, which includes the effect of added mass. Based on this finding, we selected a flapping frequency range from 8 −12 Hz at 1 Hz increments with a target amplitude of around 60*°*. Actual experimental rotation amplitudes varied between 52 −58*°*; we used these measured rotation amplitudes to populate models.

Both DBET and computational models require a representation of the wing structure. For the DBET model, we use analytical expressions to calculate the wing’s natural frequencies and mode shapes [43]. We retained only a single vibration mode. For the computational model, we used Abaqus FEA. The wing was assumed to be isotropic and homogeneous. It was discretized into 2560 eight node brick elements (4 through thickness, 640 over the surface), which was sufficient for the wing’s first three natural frequencies to converge (Fig. S1). The first natural frequency occurred at 31.6 Hz and corresponded to the bending mode (Fig. 4). Prior wing characterization *in vacuo* suggests added mass will reduce this natural frequency by about 1.2 Hz, bringing it into close agreement with the experimentally measured value [33]. Note that natural frequencies calculated via FEA are typically higher than those calculated by analytically; we therefore used different Young’s modulus to ensure both computational and DBET models have the same wing natural frequencies (Tab. 1).

**Figure 4.**
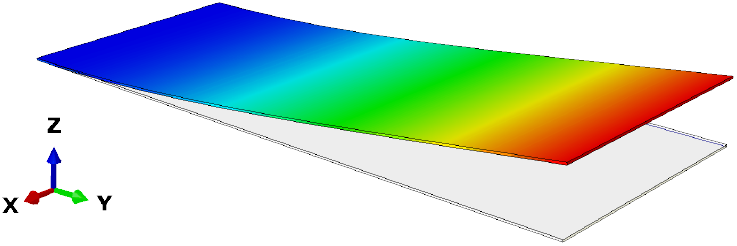
First vibration mode of the wing superimposed on undeformed wing geometry.

The background mesh region of the CFD model was sized to enclose the wing’s full flapping motion and the major flow structures leaving the wing’s surface. The overset mesh region around the wing was sized to enclose the total wing deformation and included additional clearance to minimize the creation of malformed or warped elements during the RBF morphing step. In order to increase the accuracy of the pressure calculation on the surface of the wing, prism-layers at a finer resolution were added with the default growth rate of 1.5. An appropriate element size was determined by monitoring the maximum aerodynamic moment of the wing. The average element length was decreased while ensuring that cells along the boundary of the overset mesh were never smaller than half those of the neighboring background mesh cells to minimize numerical diffusion across the Chimera boundary. Results of the mesh convergence can be seen in (Fig. S2), with a total number of elements of above one million being considered sufficiently converged relative to the computational time required. After conducting the mesh convergence, a time-step convergence study was also performed, resulting in a time-step of 1 ms, and seen in (Fig. S2). Monitoring the aerodynamic moment and tip deflection, a total of 0.5 sec were simulated to ensure the results were periodically steady.

## 4. Results

In this section, we investigate wingtip deflection measured experimentally and compare it to that calculated via DBET and computational FSI models. First, consider the experimental results. Wingtip deflection is shown as a function of stroke phase for flapping frequencies between 8-12 Hz in Fig. 5. Peak wingtip deflections range from 0.69 cm at 8 Hz to 1.83 cm at 12 Hz, representing deflections with magnitudes of 15 −37% of the wing length. For all frequencies, maximum peak deflection is delayed from the transition between the upstroke and downstroke. The delay between stroke transition and peak deflection increases with flap frequency. Because inertial forces are proportional to the angular acceleration, this delay likely results from aerodynamic effects. The wing deforms primarily at the flapping frequency, but also experiences a lesser response at three times the flapping frequency. The displacement waveform qualitatively changes with flapping frequency, suggesting the third harmonic response is sensitive to flap frequency and does not scale proportionally to the first harmonic response.

**Figure 5.**
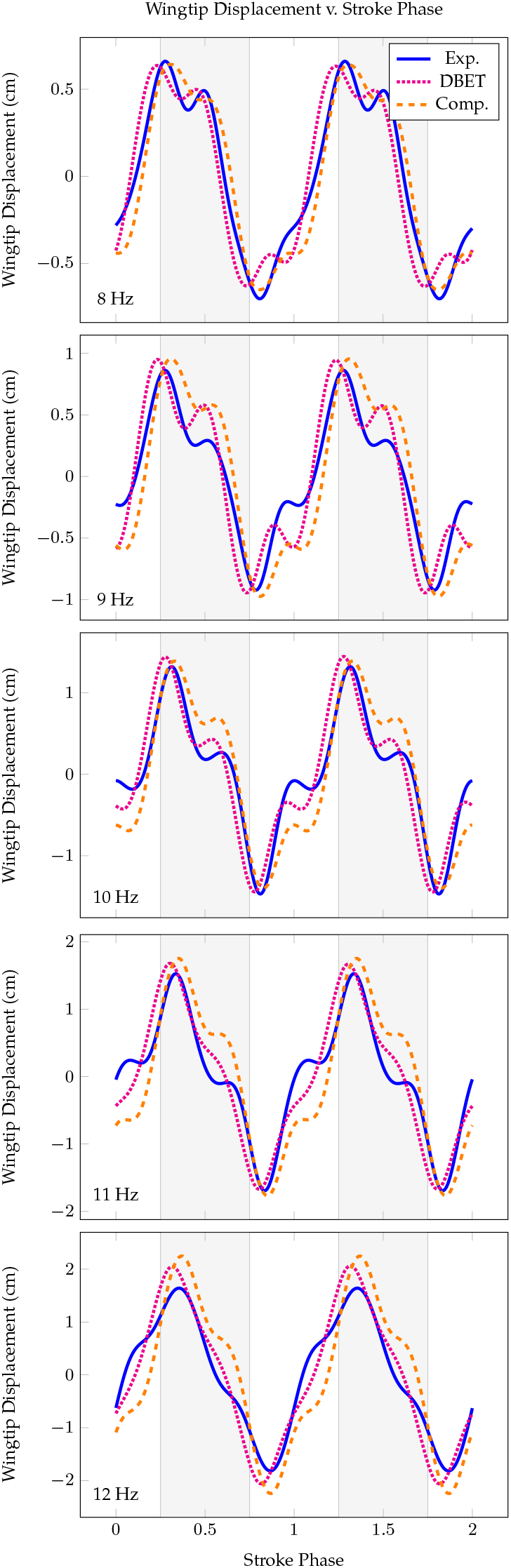
Wingtip displacements calculated via DBET and computational models are compared to experimental measurements for flapping frequencies 8 −12 Hz. Displacements are shown as a function of stroke phase over two flapping cycles. Gray regions indicate downstroke and white regions indicate upstroke.

To quantitatively compare model predictions to experimental findings, we curve fitted steady-state wingtip displacements using a fifth-order Fourier series considering three subsequent wingbeats for experimental measurements and the final wingbeat simulated for DBET and computational models. From the curve-fitted data, we calculated the overall wingtip displacement and first and third harmonic magnitudes as a function of flap frequency (Fig. 6). No other harmonics contributed significantly to wing deformation. Note that for the experimental case, overall wingtip displacement varied modestly between the upstroke and downstroke (Fig. 5). This occurred due to a weight imbalance of the wing clamping mechanism, which caused the prescribed rotation to overshoot slightly more on the downstroke than on the upstroke. We averaged maximum displacements on the upstroke and downstroke for the experimental case in order to make better comparisons against the models.

**Figure 6.**
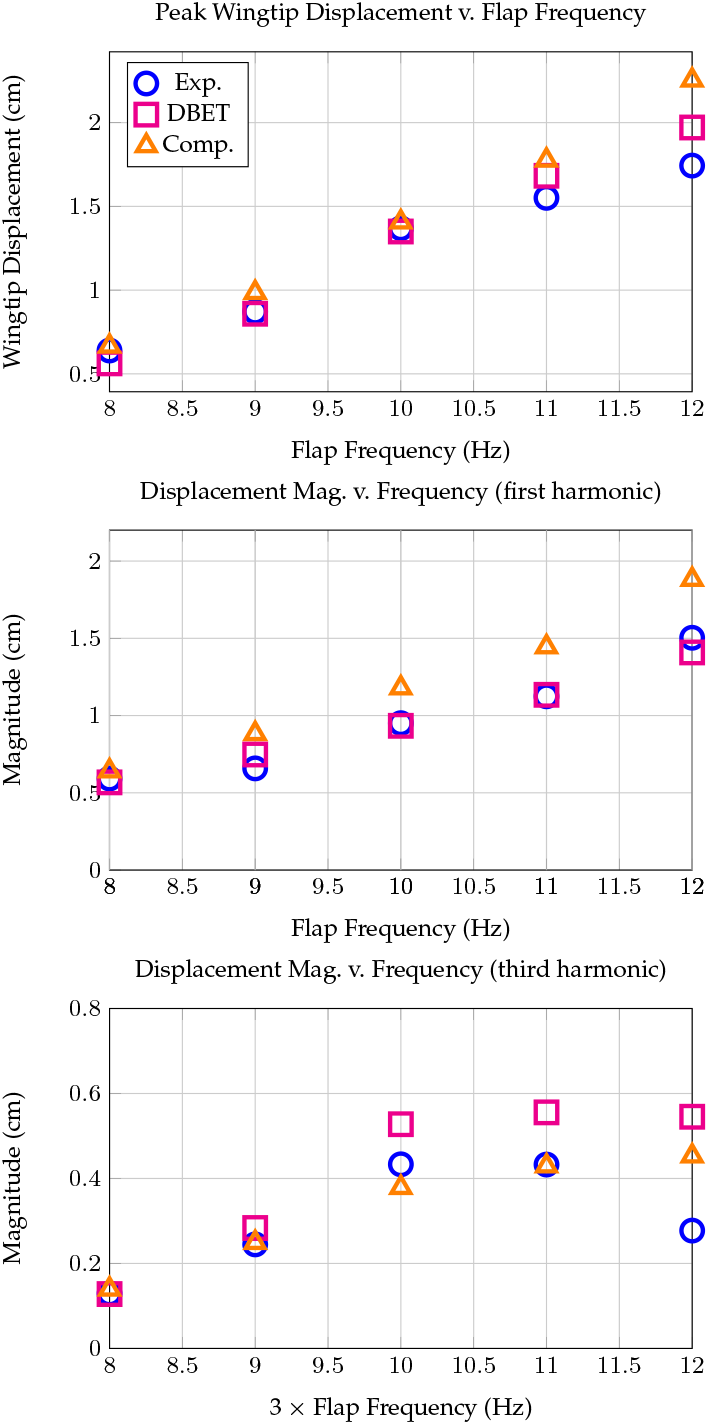
Peak wingtip displacement, as well as first and third harmonic magnitude components of wingtip displacement, as a function of flap frequency measured experimentally and as predicted by DBET and computational models.

DBET and computational models predicted the wing response fairly well. Across the considered frequency range, the DBET and computational models captured peak displacements within 13% and 30%, respectively. In agreement with the experiment, both models showed that the wing deformed at its flap frequency and three times its flap frequency. The DBET and computational models had maximum magnitude errors of 16% and 34% at the first harmonic and 72% and 63% at the third harmonic. The largest magnitude errors occur at the highest flapping frequency considered, while the error at reduced frequencies was much lower for both models.

Both models captured qualitative changes in the wing response. Each predicted that the first harmonic magnitude grows monotonically with respect to flap frequency. The experiment showed that the third harmonic magnitude experiences a local maximum and subsequently decreases. The peak in the third harmonic response indicates resonance, where the resonance condition may occur from an aerodynamic force coinciding with the wing’s first natural frequency. Previous work suggests this resonance may occur even in a vacuum due to the interaction of inertial forces with the wing’s periodically varying stiffness. Within the flapping frequency range considered, the DBET and computational model predicted a monotonic increase of the third harmonic magnitude. However, the slope of the third harmonic response suggests that its magnitude may decrease at flap frequencies exceeding 12 Hz. In general, the computational model captures the timing of the deformation waveform more accurately relative to the DBET model.

## 5. Discussion

Experimental results suggest that both the DBET and computational models are accurate within a range of flapping-to-natural frequency ratios and wingtip displacements relevant to larger flying insects. The DBET model solves for wing deformation faster than the computational model; using a workstation with 32GB of DDR4 RAM, an Intel i9-9900K 3.6GHz CPU (4 processing threads in the CFD model), the DBET model solves in about 30 ms per wingbeat whereas the computational model solves in about 8.9*×*10^6^ ms per wingbeat. This represents a 5 order of magnitude difference between the solution times achievable by DBET and computational methods. Given the level of accuracy achieved, the DBET framework is therefore a promising development that could underpin more complex flapping wing FSI models.

While economical, the DBET model has limitations that may make computational modeling desirable in some contexts. First, the computational model resolves the entire flow structure surrounding the deforming wing whereas the DBET model does not. As a result, computational models are more appropriate for capturing certain fluid phenomena such as clap-and-fling or vortex shedding. Second, the 3D computational model can capture pressure variations along the chord width. Indeed, pressure varied along the wing chord throughout the stroke cycle, though these pressures were symmetric about the wing’s centerline and did not cause it to twist. Third, the DBET framework does not account for in-plane stretching or geometric nonlinearity, though experimental results do not indicate these factors contribute significantly to the wing’s structural response. Lastly, while DBET provides a significant tool in the analysis of flapping wings, more complex fluid dynamics could be incorporated to improve the accuracy of these models, particularly as they are scaled to accommodate MDOF kinematics. Because the computational models can calculate the full fluid dynamics of a system, they will allow for better tuning of parameters that can account for phenomena, such as dynamic stall, in the future.

Despite these limitations, the semi-analytical nature of the DBET model allows us to interpret it in order to better understand the physics governing wing deformation. For example, DBET can be used to evaluate the forces acting on the wing and how these forces scale relative to one another. Understanding this physical scaling can elucidate the influence wing deformation has on different flying insects, particularly those with variable wing lengths or aspect ratios and flapping amplitudes. Derived expressions for inertial and aerodynamic forces can also be used to better understand how deformation affects other physics relevant to flight, such as power expenditures.

In what follows, we use the DBET model to investigate the forces responsible for wing deformation. The forces acting on the experimental wing flapping at 10 Hz, estimated via the DBET model, are shown as a function of stroke phase in Fig. 7.

**Figure 7.**
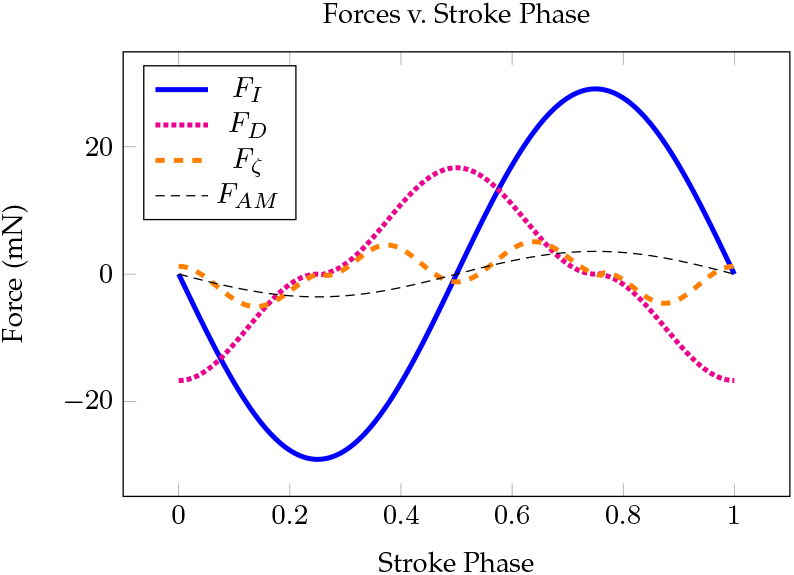
DBET model predictions of the steady-state inertial and aerodynamic forces acting on the experimental wing flapping at 10 Hz. Added mass stiffness and inertia are small compared to the forces pictured and are omitted for clarity.

### (a) Aerodynamic Drag vs. Inertia

Research suggests that deformation in Hawkmoth *M. sexta* wings, which are of comparable size to the experimental wing, is dominated by inertial forces [16]. Rigid body drag was the largest aerodynamic force acting on the wing during our experiment. The magnitude ratio between rigid body drag force *F*_*D*_ and inertial force *F*_*I*_ scales with

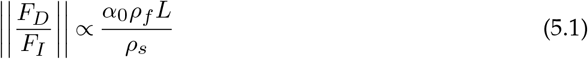

where *ρ*_*s*_ is the wing’s mass per unit area and *α*_0_ is the flapping amplitude. Intuitively, deformation will be dominated by inertial forces in heavy wings and aerodynamic forces in lightweight wings. Inertial forces increase linearly with wing length, while aerodynamic forces scale quadratically. Consequently, longer wings experience greater aerodynamic loading. When rotation amplitudes are much less than one radian, inertial forces tend to overwhelm aerodynamics. Specific to our wing and flapping amplitude, *F*_*I*_ is approximately two times greater than *F*_*D*_ (Fig. 7). However, the surface density of the paper wing is about 4-5 times larger than that of an *M. sexta* wing of comparable size. Adjusting the wing’s surface density to 20% its original value inverts this trend, where *F*_*D*_ now exceeds *F*_*I*_ by 2.5 times. Based on this simplified analysis, we conjecture that if the complex wing structure and flapping kinematics were taken into account, the aerodynamic and inertial forces contributing to *M. sexta* wing deformation are similar in magnitude.

### (b) Aerodynamic Drag vs. Aerodynamic Damping

Though rigid body drag dominates the remainder of the aerodynamic forces in most cases, there are instances where aerodynamic damping is appreciable. If the wing’s elastic response occurs primarily at the flapping frequency, the ratio between rigid body drag *F*_*D*_ and aerodynamic damping *F*_*ζ*_ is proportional to

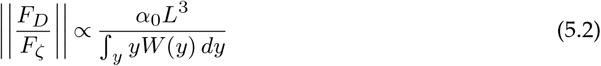

If we assume the deformed shape is a bending mode represented by 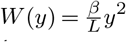, where *β* is the ratio between the wingtip deflection and wing length, this simplifies to

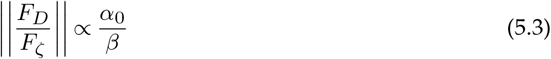

This expression demonstrates the significance of the relative magnitudes of the wing’s rigid body motion and elastic deformation. For a fixed rotation amplitude of 1 radian, aerodynamic damping will exceed rigid body drag only when wingtip deflection exceeds the length of the wing, a condition which is physically unrealistic in flight. However, for a much smaller rotation amplitude, the wingtip deflection could easily exceed 10% of wing length if the wing was highly flexible. For the experimental parameters in this work, *β* ranges about 0.15 −0.37, where rigid body drag is larger than aerodynamic damping but on the same order of magnitude.

This scaling behavior between rigid body drag and aerodynamic damping has interesting implications at resonance. Most insects flap below the first natural frequency of their wings [44], but superharmonic resonance, or resonance induced by higher-order harmonics of aerodynamic forces, may occur if the insect flaps at an integer quotient of its wing’s natural frequency. Indeed, prior work suggests flapping at 1/3 of the wing’s natural frequency may confer energetic and aerodynamic benefits [40,41]. However, these benefits would diminish if wing deformation becomes excessive. The above scaling suggests that aerodynamic damping may restrict unfavorable levels of deformation, since *β* becomes large in these contexts. In our own experiment, we found that aerodynamic damping overwhelms viscous damping of the wing structure. Thus, aerodynamic damping may play a more substantial role in attenuating large deformations relative to viscous damping inherent to the structure.

### (c) Aerodynamic Drag vs. Added Mass

After rigid body drag and aerodynamic damping, rigid body added mass is the last fluid forcing term of significance at these length scales (Fig. 7). The magnitude ratio between rigid body drag *F*_*D*_ and rigid body added mass *F*_*AM*_ is proportional to

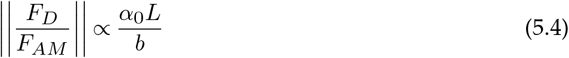

Rigid body added mass is greater in short, fat wings, whereas rigid body drag is greater in longer, more slender wings. Because rigid body added mass scales linearly with length and rigid body drag quadratically, wings with low rotation amplitudes are more influenced by the former. Added mass may also drastically lower the natural frequency of insect wings. Modal analysis in air and *in vacuo* shows that added mass reduces the first natural frequency of the *M. sexta* forewing by 30% [45]. The wing’s effective k^*th*^ natural frequency *ω*_*eff,k*_ satisfies

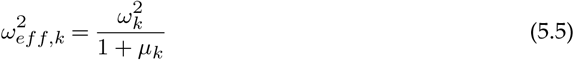

where for a cantilever beam, *µ*_*k*_ is proportional to

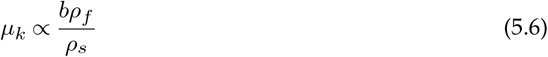

The reduction of the wing’s fundamental frequency from added mass is therefore sensitive to the ratio between the density of air and the wing’s surface density. This explains why the fundamental frequency of the *M. sexta* forewing is more greatly influenced by added mass compared to the paper wing used in our experiment.

## Supporting information

Video of mechanical flapper at 10 Hz

## Acknowledgment

This research was supported the National Science Foundation under awards Nos. CBET-1855383 to MJ and EJ and CMMI-1942810 to MJ. Any opinions, findings, and conclusions or recommendations expressed in this material are those of the author(s) and do not necessarily reflect the views of the National Science Foundation.

## Supplementary Material

**Figure S1.**
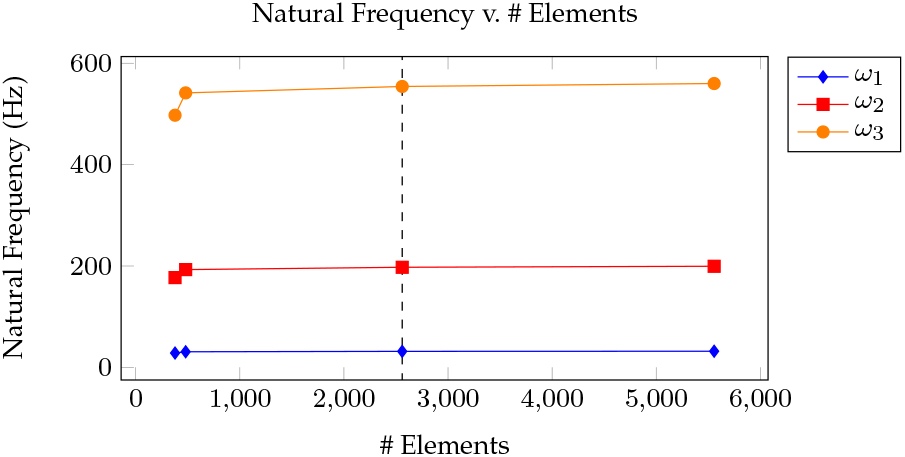
Convergence studies for the FEA model show that the first three natural frequencies converge around 2500 elements. Operating point shown in dashed lines.

**Figure S2.**
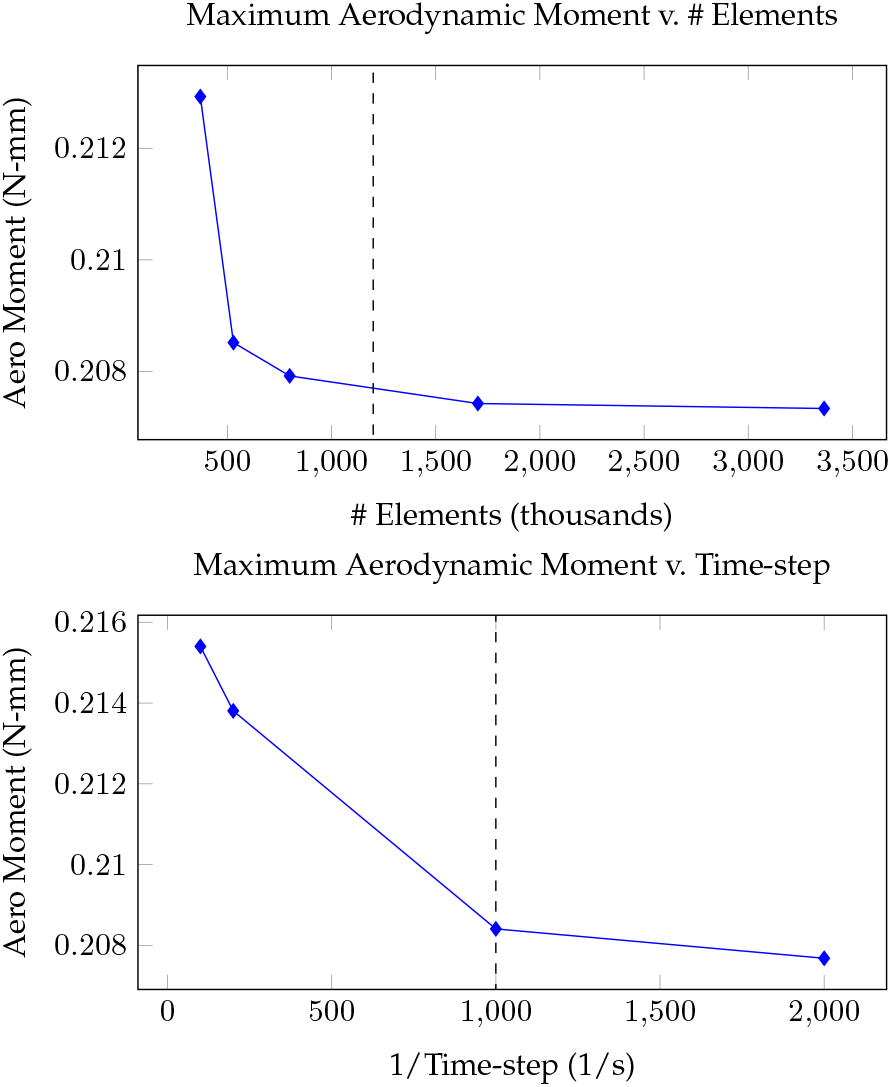
Convergence studies for the CFD model. (Top) Maximum aerodynamic moment at steady-state as a function of element count, and (Bottom) Maximum aerodynamic moment at steady-state as a function of the inverse of the time step. Operating points shown in dashed lines.

